# Topological data analysis reveals rigid brain-state dynamics during self-viewing in trait rumination

**DOI:** 10.64898/2025.12.27.696696

**Authors:** Caleb Geniesse, Sahar Jahanikia, Hua Xie, Neeraj S. Sonalkar, Leanne M. Williams, Manish Saggar

## Abstract

Rumination—repetitive, negatively valenced, self-focused thought—is a maladaptive cognitive style linked to emotional dysregulation and psychiatric risk. To investigate its neural underpinnings in a naturalistic context, we developed an fMRI paradigm in which participants observed and reflected on videos of their own past group-based problem-solving sessions, a naturalistic self-relevant context rarely examined in fMRI studies. Thirty-two adults (mean age = 30.4 ± 5.4 years; 13 F) were recorded during collaborative design-thinking tasks in triads. In a subsequent scanning session, each participant viewed two self-relevant team videos and one control team video, followed by a structured reflection period. We assessed trait rumination using the Rumination–Reflection Questionnaire (RRQ) and applied Topological Data Analysis (TDA) via the Mapper algorithm to model individual-level whole-brain dynamics during the task. Mapper shape graphs captured temporal transitions between brain states, allowing us to quantify the similarity of timepoints across the session. Individuals with higher trait rumination showed significantly higher temporal similarity, indicating reduced brain-state variability, during self-relevant conditions (r = 0.46, p = 0.018). This effect was not observed during the control condition. These findings suggest that rumination is associated with rigid brain dynamics during self-observation and evaluative processing. Traditional GLM and inter-subject correlation (ISC) analyses confirmed task engagement of key self-referential and social-evaluative regions, while Mapper revealed dynamic features not captured by static or group-averaged methods. Together, these findings demonstrate that trait rumination is associated with rigid large-scale brain dynamics during self-relevant cognition and highlight the value of combining naturalistic paradigms with topological approaches to capture behaviorally meaningful signatures.

**Significance Statement:** Rumination—repetitive, self-focused thought—is a core cognitive feature of depression and anxiety. However, the neural dynamics underlying this mental inflexibility remain poorly understood, especially during ecologically valid, self-relevant experiences. Using topological data analysis and a naturalistic self-reflection fMRI paradigm, we identify a novel neural marker: individuals with higher rumination traits exhibit reduced flexibility in brain state transitions when watching videos of themselves. This signature is uniquely captured by Mapper, a data-driven geometric framework, and not detected by traditional neuroimaging methods. Our findings offer a new window into how maladaptive thought patterns manifest in brain dynamics, paving the way for personalized diagnostics and interventions in mood disorders.

## 1. Introduction

Reflection and rumination are core components of self-focused cognition and have been widely studied across domains such as learning, creativity, and psychiatry (Christoff et al., 2009; Nolen-Hoeksema et al., 2008; Trapnell & Campbell, 1999). Reflection generally involves deliberate and adaptive self-examination, whereas rumination is characterized by repetitive, negatively valenced thoughts about one’s affective state (Treynor et al., 2003). Despite their conceptual overlap, these processes diverge in psychological outcomes: reflection is linked to insight and cognitive flexibility, whereas rumination is associated with emotional dysregulation, cognitive rigidity, and increased psychiatric vulnerability (Lyubomirsky & Nolen-Hoeksema, 1995; Whitmer & Gotlib, 2013).

It can be argued that efforts to understand the neural underpinnings of reflection and rumination should increasingly embrace naturalistic paradigms, as such ecologically valid stimuli are gaining traction for investigating cognition, personality traits, and their neural correlates (Finn et al., 2018; Matusz et al., 2019; Meer et al., 2020; Nastase et al., 2020; Saarimäki, 2021; Sonkusare et al., 2019). Traditional task-based fMRI paradigms, with controlled stimuli and rigid timing, may constrain or obscure the dynamics of spontaneous, self-directed cognition. In contrast, recent studies have leveraged naturalistic stimuli, such as films, narratives, or real-life social interactions, to better approximate the complexity of everyday cognition (Nastase et al., 2020). Analytical methods, including inter-subject correlation (ISC), representational similarity analysis, and hyperscanning, have significantly advanced our understanding of how the brain responds to dynamic social and affective contexts (Montague et al., 2002; Nastase et al., 2019; Song et al., 2021; Xie et al., 2020).

Despite this progress, important gaps remain. For example, relatively few studies employ **self-relevant stimuli**, such as personalized video or audio recordings, to probe self-referential thought (Moran et al., 2013; Northoff et al., 2006). Such personalized stimuli may provide a powerful lens into the neural mechanisms of introspection and self-evaluation. Second, many analytical approaches rely on averaging across participants, potentially masking individual variability that could be critical for understanding traits such as rumination. Third, the spatial and temporal organization and flexibility of large-scale brain dynamics, particularly during naturalistic self-relevant cognition, remain underexplored (Vidaurre et al., 2017).

To address these limitations, we developed a novel fMRI task in which participants viewed videos of themselves engaged in prior group-based problem-solving interactions. Participants worked in teams of three to solve design-thinking challenges, creating a socially embedded, ecologically valid context for self-observation. By prompting participants to critically evaluate team performance while observing themselves, we aimed to elicit reflective and potentially ruminative thought processes in a minimally structured manner. We refer to this as a self-video fMRI paradigm, designed to evoke spontaneous self-focused cognition within a naturalistic setting.

To characterize the neural dynamics underlying this process, we applied Topological Data Analysis (TDA) via the Mapper algorithm, which enables data-driven, graph-based representations of brain state transitions over time (Saggar et al., 2022, 2018; Singh et al., 2007). This approach enabled us to move beyond static contrasts, characterizing the temporal structure and variability of whole-brain activity at the individual level. We have previously demonstrated that Mapper can effectively characterize and track fluctuations in brain activity during both resting-state (Saggar et al., 2022) and task-based paradigms (Geniesse et al., 2022; Saggar et al., 2018), without requiring spatial or temporal averaging. In the present study, we extend this framework to a novel application: examining changes in neural activity as participants engage with naturalistic self-relevant video stimuli during an fMRI paradigm.

Our investigation focused on trait rumination, as assessed by the Rumination-Reflection Questionnaire (RRQ), which distinguishes maladaptive rumination from adaptive reflection (Trapnell & Campbell, 1999). By examining the relationship between Mapper-derived features and individual differences in rumination, we tested the hypothesis that greater rumination would be associated with reduced variability in brain dynamics, consistent with cognitive rigidity. Previous work has consistently linked trait rumination to cognitive inflexibility and mental rigidity, both theoretically and behaviorally (Davis & Nolen-Hoeksema, 2000; De Lissnyder et al., 2012; Koster et al., 2011; Whitmer & Gotlib, 2013). We hypothesize that trait rumination manifests neurally as reduced flexibility in dynamic brain state transitions, consistent with cognitive rigidity.

Overall, this paper introduces our naturalistic self-video paradigm and presents an initial exploratory analysis of the resulting fMRI data. Specifically, we report: (1) validation of the paradigm using conventional analyses of task-evoked brain responses and inter-subject correlation analysis; (2) adaptation of Mapper-based TDA to extract structured, interpretable features from unstructured, naturalistic fMRI data; (3) topological and spatial characterization of the resulting brain state graphs, including transitions and distributions; and (4) quantitative associations between graph-based variability measures and individual rumination scores. Together, these contributions offer new insights into how the brain dynamically configures during self-focused evaluation, extending our understanding of neural mechanisms linked to trait rumination in healthy adults.

## 2. Methods

### 2.1 A Naturalistic fMRI Video Self-Reflection Experiment

To study the neural bases associated with trait rumination, we characterized whole-brain fMRI dynamics while individuals watched and reflected on recordings of their previous social interactions. 32 adult volunteers (age: 30.4±5.4 years, 13 female, four left-handed) were recruited (final analyzed n=27 after prespecified exclusions). They attended group brainstorming sessions where they solved three design challenges with two other participants. We used the Rumination-Reflection Questionnaire to measure trait rumination. In a follow-up session, on a different day, fMRI data were collected while individuals watched 3-minute video recordings of the brainstorming session, including two videos of their team and a third showing a control team. After each video, participants were asked to reflect on the team’s performance for 90 seconds. See Figure 1 for a graphical overview of the paradigm.

**Figure 1.**
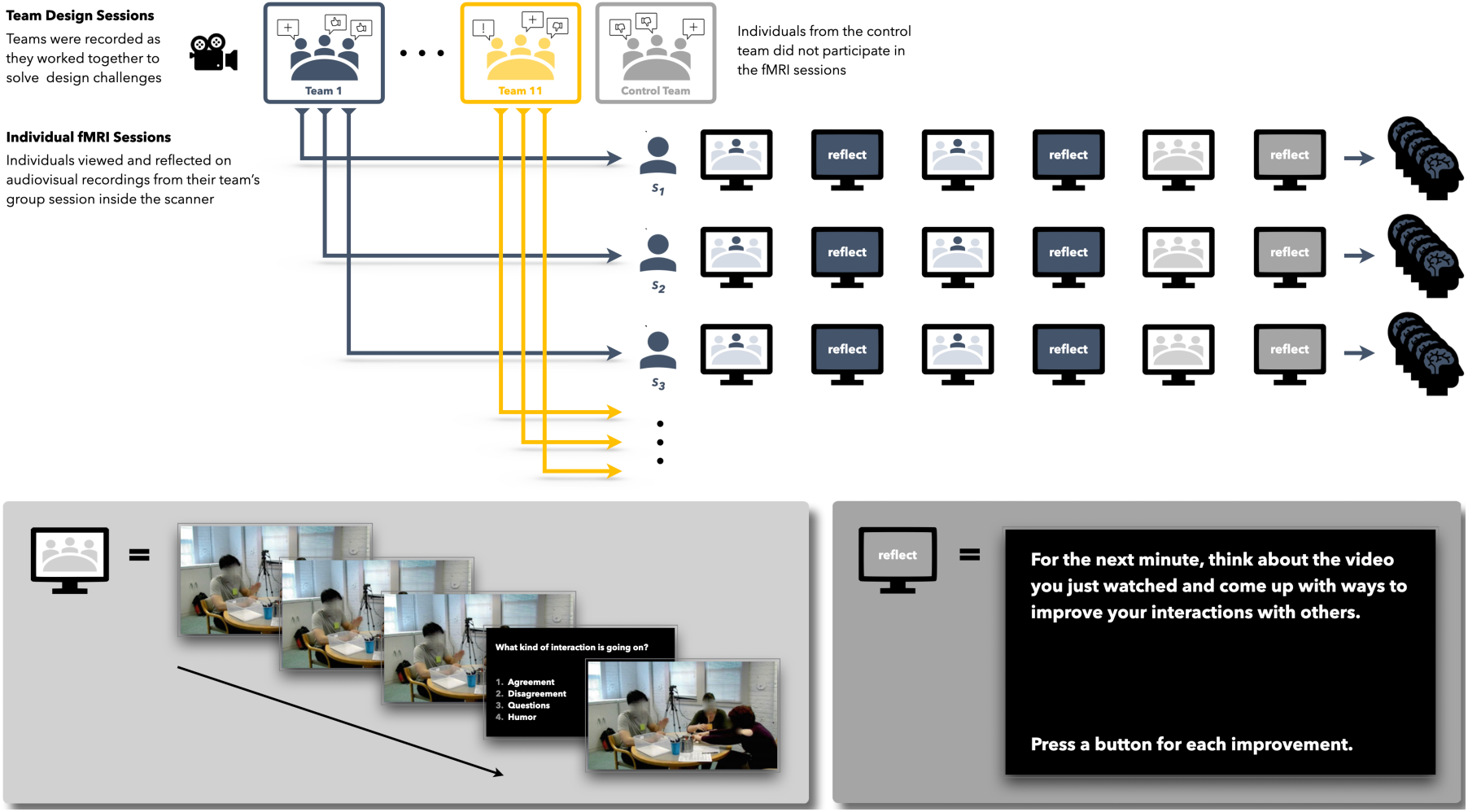
Overview of self-video fMRI paradigm: (**A**) Participants attended a group design session where they worked with two other individuals to solve various design challenges. Each design challenge included a five-minute group brainstorming session. These brainstorming sessions were recorded (audio and video) and later processed into video clips (with audio). In a follow-up fMRI scanning session, individuals viewed (and listened to) these audiovisual recordings of these brainstorming sessions inside the MRI scanner, including two of their own team (viewed only by other individuals on that team) and a third showing a control team (viewed by all participants). (**B**) Here we show example frames from the control video, including the question screen that appeared at various points during the different movies, asking participants to classify the ongoing group interaction (by pressing a button corresponding to one of four answer choices). Note, the only differences between the third control team’s video and the first two team videos (selected so that individuals viewed their own team) are the people in the videos, the design task they were brainstorming for, and possibly the camera angle. (**C**) After each video, participants were asked to reflect on the team’s performance. The question was displayed for the duration of the reflection task.

#### 2.1.1 Participants

Thirty-two participants (30.4±5.4 years, 13F, 4 left-handed) took part in our study. All participants reported no history of neurological disorder or psychotropic medication, with normal or corrected-to-normal vision. The study was approved by Stanford University’s Institutional Review Board, and all participants gave written consent.

#### 2.1.2 Behavioral Assessments

Behavioral assessments were conducted outside the MR scanner to measure trait rumination and reflection, and other standard personality traits. Here, we focus on trait rumination and reflection scores measured using the Rumination Reflection Questionnaire (RRQ) (Trapnell & Campbell, 1999). The RRQ measures a participant’s tendency to self-ruminate and self-reflect.

#### 2.1.3 Team Activity Phase

In the team activity phase, participants worked in triads to complete three rapid design challenges. Each design activity consisted of three sequential stages: a team brainstorming stage, in which participants interacted to collaboratively generate solution concepts; an individual prototyping stage, where each participant independently built a prototype using provided materials; and a team prototyping stage, where participants reconvened to synthesize their prototypes into a shared solution.

The design activities took place in the Design Observatory at the Center for Design Research at Stanford University, with video recorded from four camera angles to document the interactions. Audio was recorded separately and later synchronized with the video. To analyze communication patterns, we applied Interaction Dynamics Notation (IDN) (Sonalkar et al., 2013) to five-minute excerpts from the team brainstorming stage of each activity, as this phase contained the most interactive content.

After the IDN analysis, each brainstorming section was segmented into shorter video clips of varying durations (10–100 seconds), with each clip highlighting specific aspects of the team interaction, such as asking questions, expressing humor, or showing agreement or disagreement. These curated clips were then shown to participants during the fMRI scanning session. A separate pilot testing phase (n = 6) had revealed that focusing on the brainstorming phase was more engaging and informative for self-reflection than using the full 15-minute design activities (Sonalkar et al., 2020). Audio/video recordings captured each team’s interaction during the design activities. Participants were informed that their interactions would be recorded for research purposes, and they were asked to sign a consent form.

#### 2.1.4 Naturalistic Viewing Phase

In the fMRI experiment, participants viewed the three video clips derived from their team-activity phase, each approximately three minutes in duration. Within each clip, prompts derived from the IDN analysis were inserted, and the video was paused at predefined timepoints. Participants were then asked to identify the type of ongoing interaction, such as “agreement,” “disagreement,” “humor,” or “questioning.” This recognition task served primarily as an explicit attention check to ensure participants remained engaged with the stimuli. Following each video, participants completed a one-minute reflection phase in which they were instructed to think about the video they had just seen and come up with ways to improve their own interaction with others in similar group settings.

### 2.2 fMRI Data Acquisition and Preprocessing

#### 2.2.1 Data Acquisition

Participants were scanned using a GE 3T Discovery MR750 scanner with a 32-channel Nova Medical head-coil at the Stanford Center for Cognitive and Neurobiological Imaging. Functional scan parameters used are as follows: 1270 volumes, repetition time TR = 0.71 s, echo time TE = 30 ms; flip angle FA = 54, field of view FOV = 220 × 220 × 144 mm, isotropic voxel size = 2.4 mm, #slices = 60, multiband acceleration factor = 6. High-resolution T1-weighted structural images were also collected with FOV = 190 × 256 × 256 mm, FA = 12, TE = 2.54 ms, and isotropic voxel size = 0.9 mm.

#### 2.2.2 Data Preprocessing

We discarded the first 10 frames of each participant’s functional data to allow for T1-equilibration. Subsequently, we applied a standardized preprocessing pipeline using fMRIPrep (version 2.20.0; (Esteban et al., 2019)). This pipeline included motion correction, slice-timing correction, susceptibility distortion correction, and normalization to the Montreal Neurological Institute (MNI152) template (Evans et al., 2012).

Following fMRIPrep, additional preprocessing steps were performed in MATLAB. We discarded an additional 10 frames to account for non-stationary signal, and restricted analyses to voxels within gray matter as defined by the 3D spatial brain masks provided by fMRIPrep. Gray-matter voxel time series were extracted from the preprocessed 4D fMRI data and reshaped into a 2D matrix (voxels × time). Individual time frames with high framewise displacement (FD > 0.5 mm) were marked as motion outliers; the data for these frames were replaced with NaN values so they could be ignored in later processing.

After motion censoring, voxel-wise data were demeaned and detrended. Nuisance regression was performed for each voxel to remove physiological noise (signals from white matter and cerebrospinal fluid) and motion-related noise (six head-motion parameters: three translations and three rotations). Linear interpolation was then applied to smooth over missing data corresponding to censored high-motion frames. Finally, temporal bandpass filtering (0.009 Hz < f < 0.08 Hz) was applied.

For region-based analyses, mean time series were extracted from 375 predefined regions of interest (ROIs) based on the multi-atlas parcellation of Shine et al., (2016), which combines 333 cortical regions from the Gordon atlas (Gordon et al., 2017), 14 subcortical regions from the Harvard-Oxford subcortical atlas, and 28 cerebellar regions from the SUIT atlas (Diedrichsen et al., 2009). Prior to running further analyses, any ROIs with zero variance were excluded, and the remaining data were z-scored.

A total of 32 participants were invited to return for individual fMRI scanning on a separate day, typically one week after the initial team-based design session. Of these, two participants were excluded prior to preprocessing due to failure to complete the scanning session. An additional five participants were excluded prior to any analysis: three due to technical issues during scanning, one due to poor structural registration by fMRIPrep, and one due to excessive head motion (more than 10% of frames with FD > 0.5 mm). This resulted in a final dataset of 27 participants included in the analyses.

### 2.3 Data Analyses

#### 2.3.1 Task-Related Activation Captured by GLM Analysis

To validate the ecological validity of our naturalistic self-video paradigm, we first examined whether consistent brain activations could be detected within and across tasks as participants viewed recordings of their own team’s design sessions compared to those of a control team.

Specifically, we anchored the task to brain anatomy by applying a traditional general linear model (GLM) approach. The experimental task design was used to construct six explanatory variables corresponding to the video-viewing conditions. Within each participant, two contrasts were computed (each self-viewing video minus the control video) to identify brain regions showing stronger positive associations with self-relevant video stimuli. These within-subject contrasts were then averaged to produce a single average contrast (self-viewing minus control) for each participant.

At the group level, we conducted a non-parametric permutation regression analysis using FSL’s randomize tool to combine region-wise (rather than voxel-wise) contrasts across participants. Finally, to visualize the group-level results on the brain, the regional values were mapped back to their corresponding voxels.

#### 2.3.2 Inter-Subject Correlation (ISC) Analysis

Next, we examined regions in the brain that showed synchronized activity across participants as they viewed recordings of their own team’s design sessions, compared to a control team.

Given that participants viewed different self-relevant videos (their own team’s recordings) while the control video was identical across all participants, we reasoned that self-relevant content might evoke heterogeneous but meaningful neural responses. Some participants might show strong synchrony with certain other viewers who engaged similarly with self-relevant material, even if not all participant pairs were equally synchronized.

We implemented a variant of ISC analysis. For each brain region, we computed pairwise correlations between each participant’s time series and those of every other participant, yielding a participant-by-participant correlation matrix. To characterize individual synchrony, each participant was then assigned their maximum correlation with any other participant (i.e., the row-wise maximum of the correlation matrix). This approach departs from conventional group-level ISC methods by preserving participant-specific synchrony scores, thereby capturing individual differences. Instead of aggregating a single ISC value across all participants, we retained a loop over each participant to compute a statistic (e.g., mean or maximum) based on that participant’s pairwise correlations with the remainder of the group. This approach was also motivated by practical constraints, as some teams had only two members with usable fMRI data, precluding within-team ISC calculations for those groups.

As in the GLM analysis, the experimental task design was used to define six explanatory variables. Within each participant, two contrasts were computed (each self-viewing video minus the control video) to identify brain regions showing stronger positive synchrony with self-relevant videos. These contrasts were then averaged within each participant to yield a single self-viewing-minus-control contrast score. At the group level, a non-parametric permutation regression analysis using FSL’s randomize tool was applied to combine region-wise contrasts across participants.

#### 2.3.3 Topological Data Analysis using Mapper

To capture dynamic insights about how brain activity changes over time at the level of individual participants, we applied the Mapper framework (Geniesse et al., 2022, 2019; Saggar et al., 2022, 2018; Singh et al., 2007) and developed novel analyses tailored to our naturalistic paradigm. Figure 2 provides an overview of the standard Mapper pipeline.

**Figure 2.**
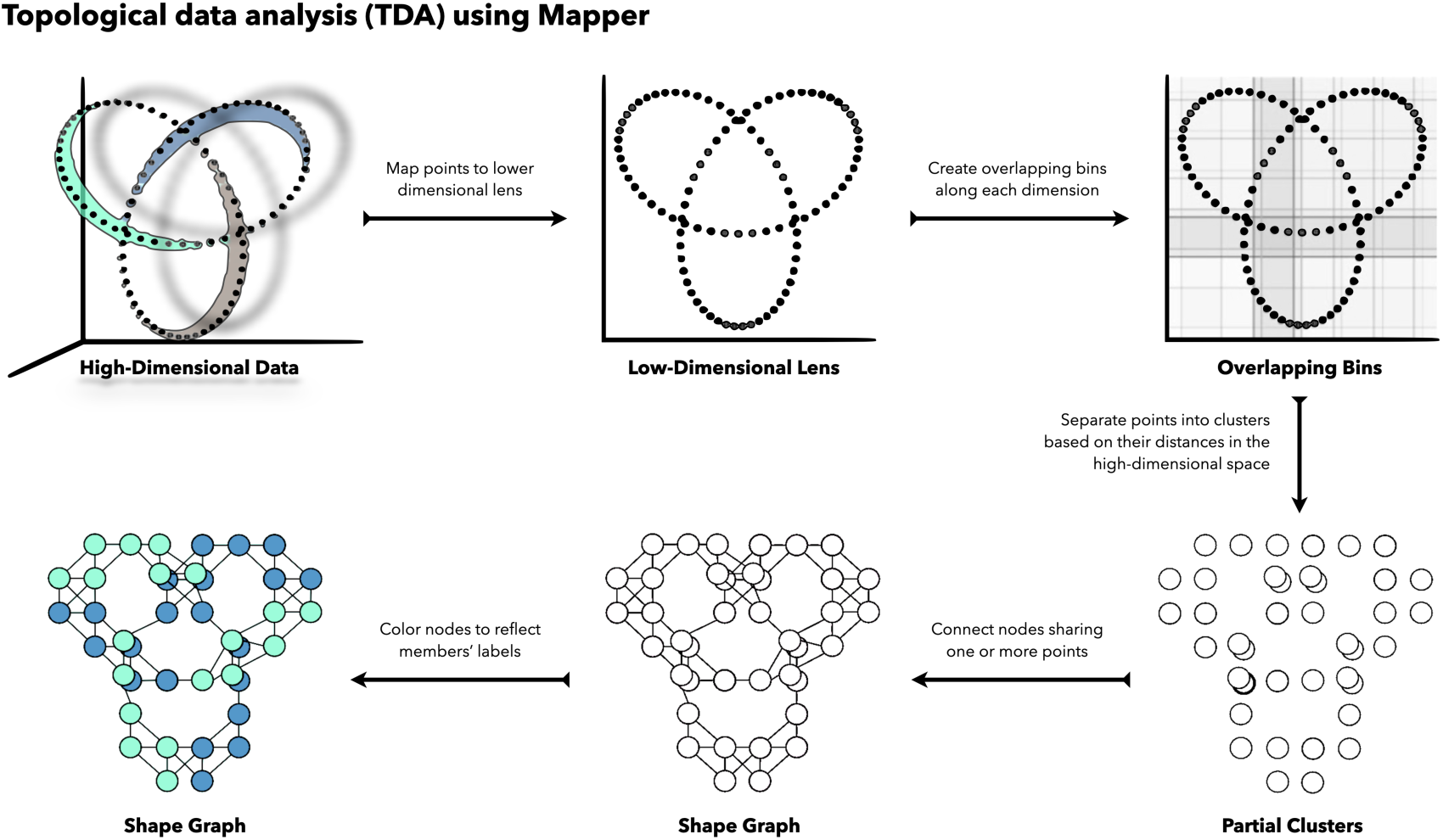
Overview of the Mapper Algorithm. Starting with a (typically high-dimensional) point cloud, and provided a lens (i.e., low-dimensional projection of the data, or a set of one or more filter functions), Mapper partitions the space into overlapping bins and maps data points into these bins. To separate points in a bin that are far apart in the high-dimensional space, Mapper performs partial clustering on the points in each bin, separately. The resulting clusters are used as nodes in the final graph representation. Edges are then placed between any pair of nodes associated with one or more of the same data points. The final graph representation can then be annotated to reflect node-level properties (e.g., number of edges) or data-driven variables (e.g., proportion of data points associated with each class in a set of categorical labels; average value of a function evaluated on each of the underlying data points).

Mapper requires an initial low-dimensional projection of the high-dimensional data as a “lens.” Before applying Mapper, we systematically explored both linear and non-linear dimensionality reduction techniques to identify the most suitable projection for our dataset. **Figure 3** shows five different two-dimensional projections of a representative participant’s data. Among these, Reciprocal Isomap appeared to best capture both local and global data structure, including local temporal similarity driven by the autocorrelation structure of the BOLD signal. Reciprocal Isomap improves on standard Isomap by introducing a reciprocity constraint: neighbors of a point must also reciprocally include that point as a neighbor, thereby pruning unidirectional connections.

**Figure 3.**
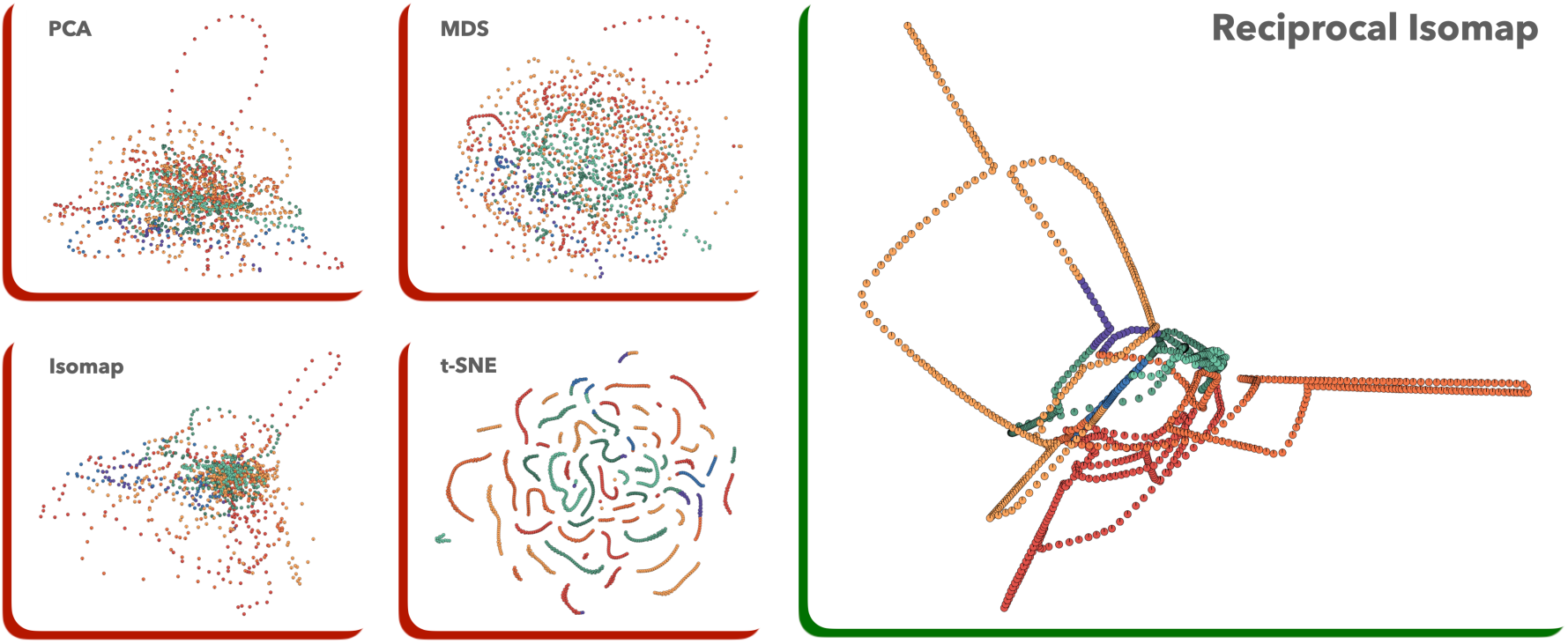
Comparing Linear and Non-Linear Embeddings of Naturalistic Data. Here, we compare different 2D embeddings of fMRI data from a representative participant (i.e., *S*_27_). On the left we show four embeddings created using linear (e.g., PCA) and non-linear (e.g., MDS, Isomap, t-SNE) methods. We consider these embeddings to be less than ideal, as they fail to capture important temporal structure in the data while also capturing higher within-task similarity as delineated by the experimental paradigm. On the right, we show an ideal embedding of the data created using our Reciprocal Isomap, a variant of the Isomap algorithm that enforces symmetric neighbor connectivity in the underlying *k*-Nearest Neighbor graph.

This approach was inspired by our previous work with NeuMapper, where we represented data directly in high-dimensional space via a k-nearest-neighbor graph, while enforcing the reciprocity condition to enhance robustness against noise and over-connection (Geniesse et al., 2022; Saggar et al., 2022). For Reciprocal Isomap, geodesic distances are computed along the resulting reciprocal graph, and these distances are then embedded into a low-dimensional projection using classical multidimensional scaling (MDS) or kernel PCA — similar in spirit to standard Isomap, but with improved graph pruning. Although similar ideas have been noted in the literature, they are not typically provided in standard machine learning libraries. Therefore, we initially implemented Reciprocal Isomap in MATLAB and subsequently developed an open-source Python implementation compatible with scikit-learn design principles (https://github.com/calebgeniesse/reciprocal_isomap).

Having established Reciprocal Isomap as a suitable projection, we used it as the lens for Mapper. We applied the Mapper algorithm as described in previous work (Haşegan et al., 2024; Saggar et al., 2022, 2018), using the L¹ (Manhattan) distance metric, which is advantageous for high-dimensional fMRI data. Mapper parameters were chosen using heuristics informed by prior studies: intermediate resolution (r = 15), gain (g = 70), and a neighborhood size of k =15, which reflects the temporal resolution of the BOLD signal. Results were stable across a wide variety of Mapper parameters (see Supplementary Figure S1). Partial clustering was performed with single-linkage hierarchical clustering, and histogram-based methods were used to identify appropriate hyperparameters.

Graph construction proceeded by treating each cluster as a node and drawing edges between nodes that shared one or more time points. Additional graph-based representations and visualizations were generated using DyNeuSR (Geniesse et al., 2019). **Figure 4** shows an example Mapper graph with uniform node coloring. It is worth noting that the layout of the graph preserves the geometry of the original embedding only for visualization purposes; nodes are positioned using the average coordinates of their members in the embedded space. Alternative layouts, such as a force-directed graph, would not preserve this geometry, but the topological relationships among nodes would remain intact. In other words, if two points are near each other in the high-dimensional space and in the embedding, they are likely assigned to the same or neighboring nodes. Conversely, if points are close in the embedding but distant in the high-dimensional space, they will typically not share the same node due to separation by the partial clustering step.

**Figure 4:**
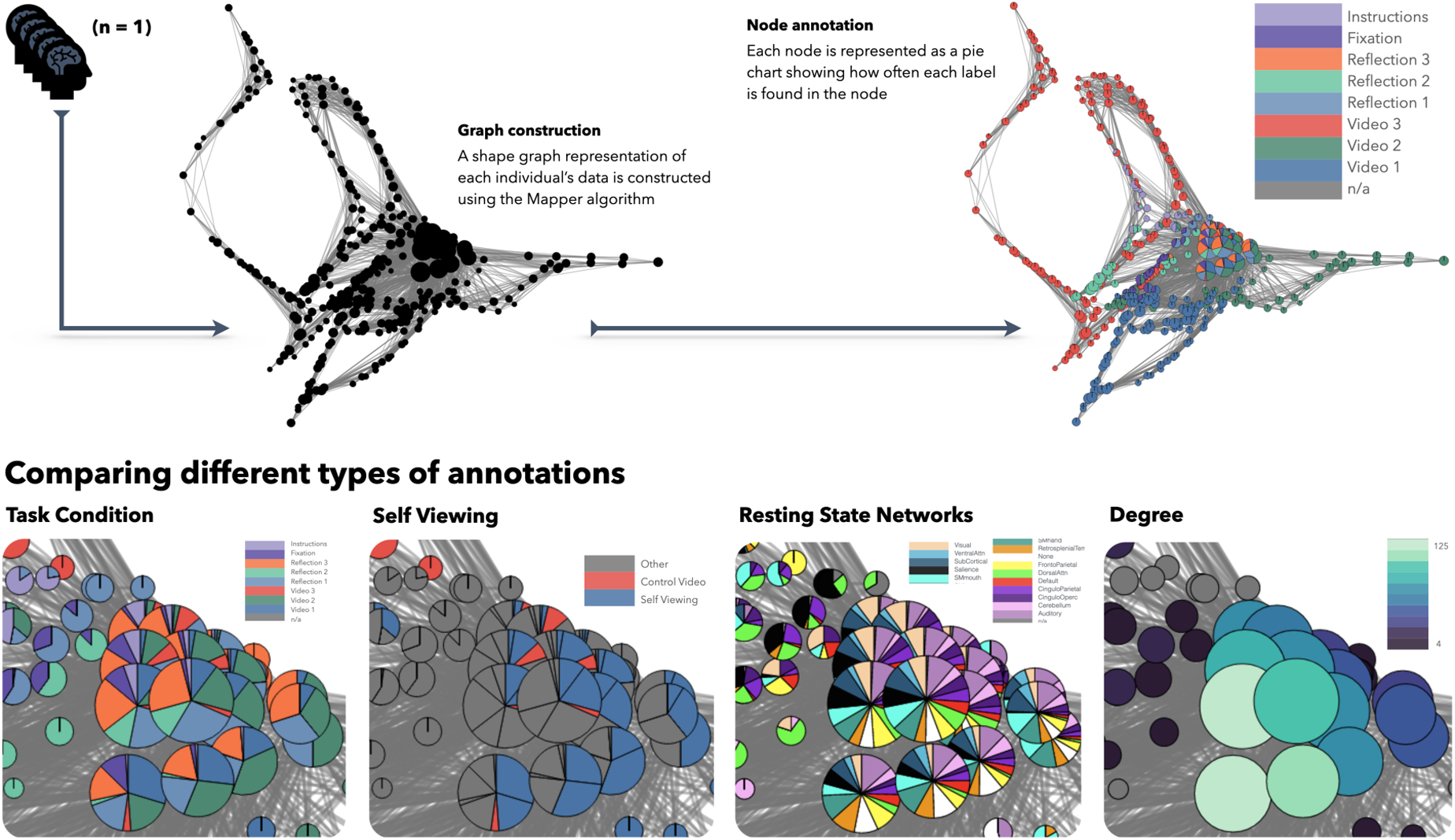
Generating and Annotating Mapper Graphs. (**A**) Shows the Mapper graph for a representative participant (i.e., *S*_27_) after initial construction, before and after annotating nodes based on the experimental task design. (**B**) We zoom into the dense central core of the graph to show how the same nodes can be annotated in a variety of ways using categorical labels and continuous values. Here, we compare annotations derived from the experimental task design, self-viewing versus control video conditions, resting-state network activations, and nodal degree values. Qualitatively, the more central and densely connected nodes have a high degree and more uniform proportions of experimental task and resting-state network annotations. In contrast, while the self-viewing condition makes up a large proportion (sometimes more than half) of most of these central nodes, the control video is rarely observed in this region of the graph (and only represents a small fraction of any central nodes it does appear in).

#### 2.3.4 Annotating and Coloring Mapper Graphs

After constructing the Mapper graphs, we leveraged the rich information provided by the naturalistic paradigm to explore diverse ways of annotating these graphs. Given categorical labels, we annotated each node by calculating the proportion of its member time points associated with each label. These proportions were then visualized using pie charts. For continuous variables, we instead assigned each node the average value of that variable across its members and represented it with a single color. Alternatively, continuous values could be discretized into bins, with each bin assigned a distinct color, to facilitate clearer interpretation.

**Figure 4** provides a zoomed-in view of an example Mapper graph, illustrating four different ways of annotating its nodes. To highlight the flexibility of our approach, we qualitatively compared several annotation schemes, capturing a range of experimental and data-driven features.

##### Annotating by experiment design variables (e.g., task conditions)

The most direct annotation approach leverages the experimental block structure. Here, we annotated nodes according to task conditions, distinguishing between individual video-viewing periods and subsequent reflection periods. We also created a combined annotation that grouped all video segments together versus all reflection periods together, providing a simplified contrast between these task epochs. We created additional condition design variables by combining the design variables for tasks assigned to each condition. For example, to create a self-viewing condition variable, we combined the design variables for videos 1 and 2. We did this for various conditions, including self-viewing, self-reflection (i.e., self-viewing videos), viewing (i.e., self-viewing and control videos), reflection (i.e., self and control reflection tasks), self (i.e., self-viewing videos and their subsequent reflection tasks), control (i.e., control video and reflection).

##### Annotating by activation in resting state networks (RSN)

Next, we derived data-driven annotations based on functional networks. For each time point, we averaged activation across parcels within canonical resting-state networks (RSNs). We then z-scored these averages and masked activations below one standard deviation above the mean, allowing each time point to potentially belong to multiple active networks. A node’s pie chart thus represented the frequency with which each network was active across its member time points. We generated framewise annotations describing the relative engagement of 14 canonical resting-state networks (including subcortical, cerebellum, and a “None” label corresponding to regions not assigned to any of the networks). Here, we considered a network to be active if the mean activity (averaged across parcels in the network) was greater than one standard deviation above the mean (averaged across all networks).

##### Annotating by topological properties of the graph (e.g., node degree)

We first considered annotations that rely solely on the graph’s topological properties, without requiring any external information. Beyond uniform node coloring, nodes can be colored according to topological metrics such as node degree. Examining degree-based annotations, for example, reveals that high-degree nodes often contain a greater diversity of member time points, which can be reflected in a more complex pie-chart annotation.

#### 2.3.5 Characterizing Mapper Graphs

To extract insights about brain dynamics from individual Mapper graphs at the level of single time frames, we constructed a temporal similarity matrix (TSM) for each participant. The TSM captures how time frames are related to each other within the Mapper graph based on their co-occurrence within graph nodes.

Specifically, we computed a temporal co-occurrence matrix (COO), which counts how often pairs of time frames are assigned to the same node. The COO can be efficiently calculated as the dot product of the membership matrix with its own transpose.

Having computed the TSM for each participant, we then quantified its topological properties using network science techniques. For each time frame, we calculated its temporal similarity degree as the row-wise average of the TSM, reflecting how strongly that time frame is connected to other time frames. To normalize these values, we divided each degree by the maximum degree across all time frames for that participant.

Finally, to relate these temporal similarity features to trait rumination, we computed the Pearson correlation coefficient between the normalized temporal similarity degree and participants’ rumination scores on the RRQ.

## 3. Results

We sought to validate our novel, ecologically grounded fMRI paradigm, based on participants watching and reflecting on real-world group interactions, by applying multiple complementary analysis strategies. First, we used traditional task-based and naturalistic analysis methods to extract group-level effects: (1) general linear modeling (GLM) to identify condition-specific activation within individuals, and (2) inter-subject correlation (ISC) to capture shared neural synchrony across participants. These analyses anchored our paradigm in existing neural frameworks and informed our interpretation of individual-level topological dynamics. Finally, we applied Topological Data Analysis (TDA) using the Mapper algorithm to examine brain state transitions and their associations with trait rumination scores.

### 3.1 Enhanced Self-Referential Activation During Self-Viewing: Evidence from GLM Analysis

We first examined task-related activation using a general linear model (GLM) framework. Specifically, we tested whether viewing videos of oneself in a real-world group interaction would evoke distinct patterns of neural activation compared to viewing a control team. At the first level, subject-specific GLMs were constructed using six explanatory variables derived from the task design. Within each subject, two primary contrasts were generated: (1) self-viewing video greater than control video, and (2) self-reflection greater than control reflection. These contrast maps were then entered into group-level analysis using FSL’s *randomise* tool with threshold-free cluster enhancement (TFCE), allowing us to identify regions showing consistent activation differences across participants.

For the self-viewing greater than control video contrast, we observed significant clusters of increased activation in the left anterior cingulate cortex (ACC) and paracingulate gyrus, extending dorsally into the supplementary motor area (SMA). Additional activation was found in the right lateral occipital cortex (inferior division), bilateral inferior temporal gyri, bilateral middle temporal gyri, and bilateral temporal-occipital fusiform cortex (**Figure 5**). These regions are broadly associated with self-referential processing, social cognition, and complex visual analysis, supporting the idea that watching one’s own behavior in a group setting evokes rich cognitive and affective engagement. By contrast, the self-reflection versus control reflection contrast did not yield any clusters at corrected thresholds, suggesting that the video-viewing phase elicited more consistent and robust neural differentiation than the subsequent reflection period. These results support the sensitivity of our naturalistic task design in evoking individualized yet measurable brain responses to self-relevant social stimuli.

**Figure 5.**
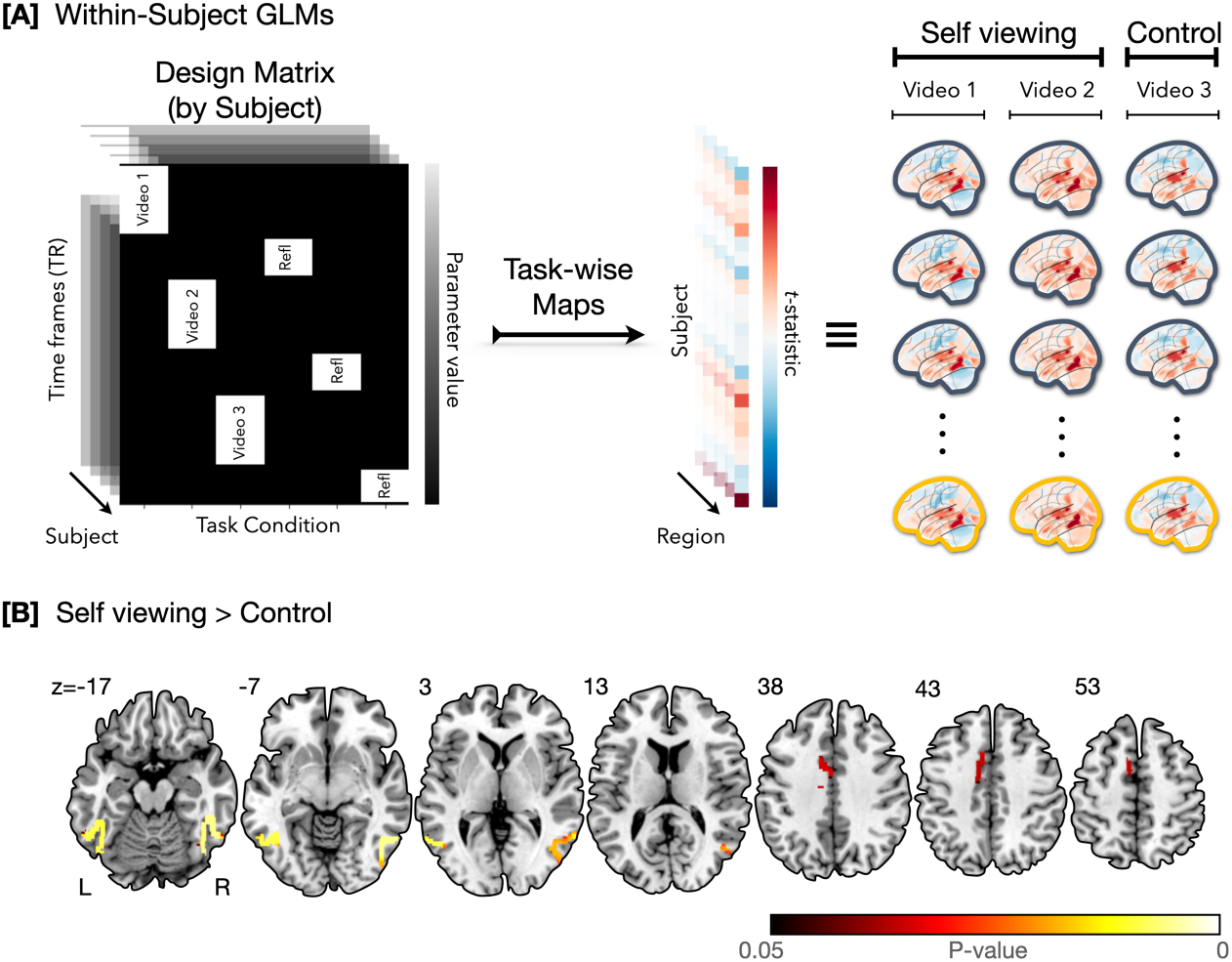
Task-based activation analysis comparing self-viewing to control videos. (**A**) Schematic of the within-subject general linear model (GLM) analysis pipeline. For each participant, we modeled brain activity associated with viewing three task conditions: two self-viewing videos and one control video, followed by brief periods of self-evaluation. Beta maps were computed for each video and each subject, and contrasts (self-viewing > control) were entered into group-level analysis. (**B**) Results of group-level GLM analysis (N=26) comparing brain responses during self-viewing videos versus the control video, using FSL’s randomise with threshold-free cluster enhancement (TFCE). Significant activation was observed in the anterior cingulate cortex, supplementary motor area, bilateral middle temporal gyrus, inferior temporal gyrus, and fusiform cortex, consistent with regions involved in self-referential and visual social processing.

We next tested whether the strength of activation in self-viewing > control correlated with individual trait rumination scores. Across all clusters derived from GLM contrasts, no significant associations with rumination were observed (all *p-*values > 0.3). This absence of trait associations contrasts with the Mapper-based results below, underscoring the sensitivity of dynamic, individual-level analyses.

### 3.2 Participant-Specific ISC Reveals Self-Viewing Synchrony in Cerebellum

To assess shared neural processing during the self-viewing task, we computed inter-subject correlation (ISC) using a participant-specific approach that captures individual differences in neural synchrony (see Methods). In brief, each participant was assigned their maximum pairwise correlation (per brain region) with all other participants, during both self- and control-viewing conditions. These scores were entered into within-subject contrasts (Self Video 1 > Control Video, and Self Video 2 > Control Video), followed by group-level statistical testing using FSL’s randomise with threshold-free cluster enhancement (TFCE).

This analysis revealed a single significant cluster located in the right cerebellar lobules I–IV, where ISC was greater during self-viewing than control viewing (**Figure 6**). Notably, this pattern emerged despite self-videos varying across teams. At the same time, the control video was identical for all participants, a design that would typically favor stimulus-locked synchrony during control viewing. This suggests that cerebellar activity during self-relevant video engagement was more temporally synchronized across individuals, possibly reflecting shared processing of embodied, self-referential content rather than stimulus-driven synchrony.

**Figure 6.**
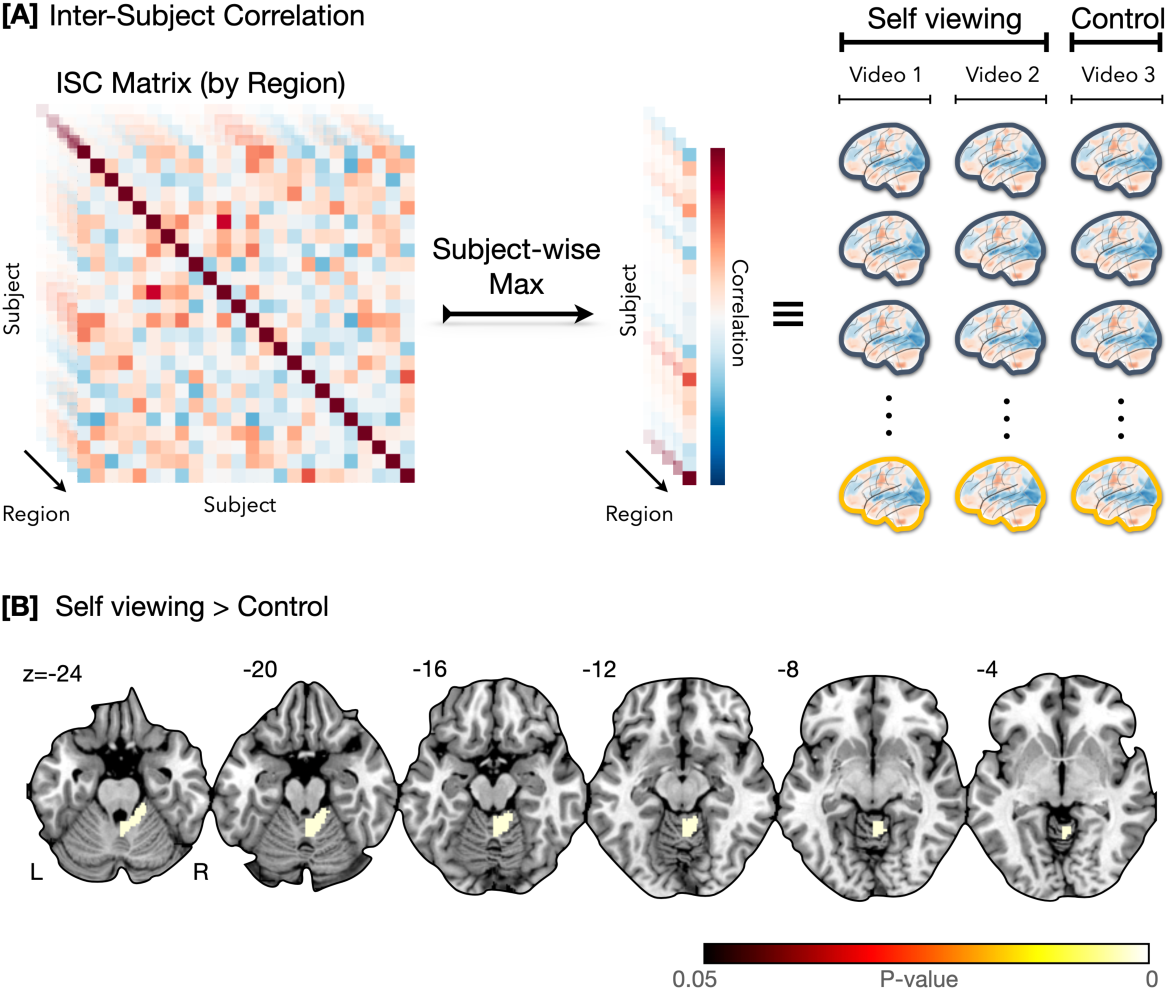
Inter-subject correlation (ISC) analysis reveals enhanced cerebellar synchrony during self-viewing. (**A**) For each participant and each brain region, we computed a subject-specific ISC score by first calculating the correlation of that participant’s time series with every other participant’s time series (ISC matrix), then selecting the maximum value per row. This yielded participant-specific ISC maps for each video, allowing us to retain individual differences in neural synchrony across conditions. (**B**) Group-level comparison of ISC maps for self-viewing (Video 1 and 2) versus control (Video 3) revealed a single significant cluster in the right cerebellum (lobules I–IV), with increased synchrony across participants during self-viewing videos. Results shown at p < 0.05 TFCE-corrected.

In contrast, when comparing the self-reflection phase to the control-reflection phase, no significant clusters were observed. This null result parallels the GLM findings and suggests that passive self-viewing, rather than reflective evaluation, drove the strongest inter-subject synchrony in this paradigm.

We next tested whether the strength of ISC in self-viewing > control correlated with individual trait rumination scores. Across all clusters derived from ISC contrasts, no significant associations with rumination were observed (all *p-*values > 0.75).

### 3.3 Mapper Reveals Increased Temporal Similarity in High Trait Ruminators

To investigate how brain dynamics during naturalistic self-reflection relate to individual differences in trait rumination, we applied the Mapper algorithm (Singh et al., 2007), a topological data analysis (TDA) technique that captures the intrinsic geometric structure of high-dimensional data. Mapper enables the construction of graphs, where each node represents a cluster of temporally adjacent fMRI timepoints with similar whole-brain activation patterns, and edges encode temporal co-occurrence. This representation retains dynamic features of the brain signal while enabling individualized visualization and quantification of brain state trajectories (Geniesse et al., 2022, 2019; Saggar et al., 2022, 2018).

For each participant, we generated a Mapper graph using their entire time series across all three video and reflection conditions. The layout of the graph reflects high-dimensional proximity between brain states, while the topology captures how those states evolve over time. As illustrated in **Figure 7A**, participants with low trait rumination (e.g., S30) showed more distributed graphs with integrated connectivity across task conditions. In contrast, high ruminators (e.g., S27) exhibited fragmented topologies where control video timepoints tended to be located in peripheral, low-degree regions, while self-relevant team videos occupied the central high-degree core.

**Figure 7.**
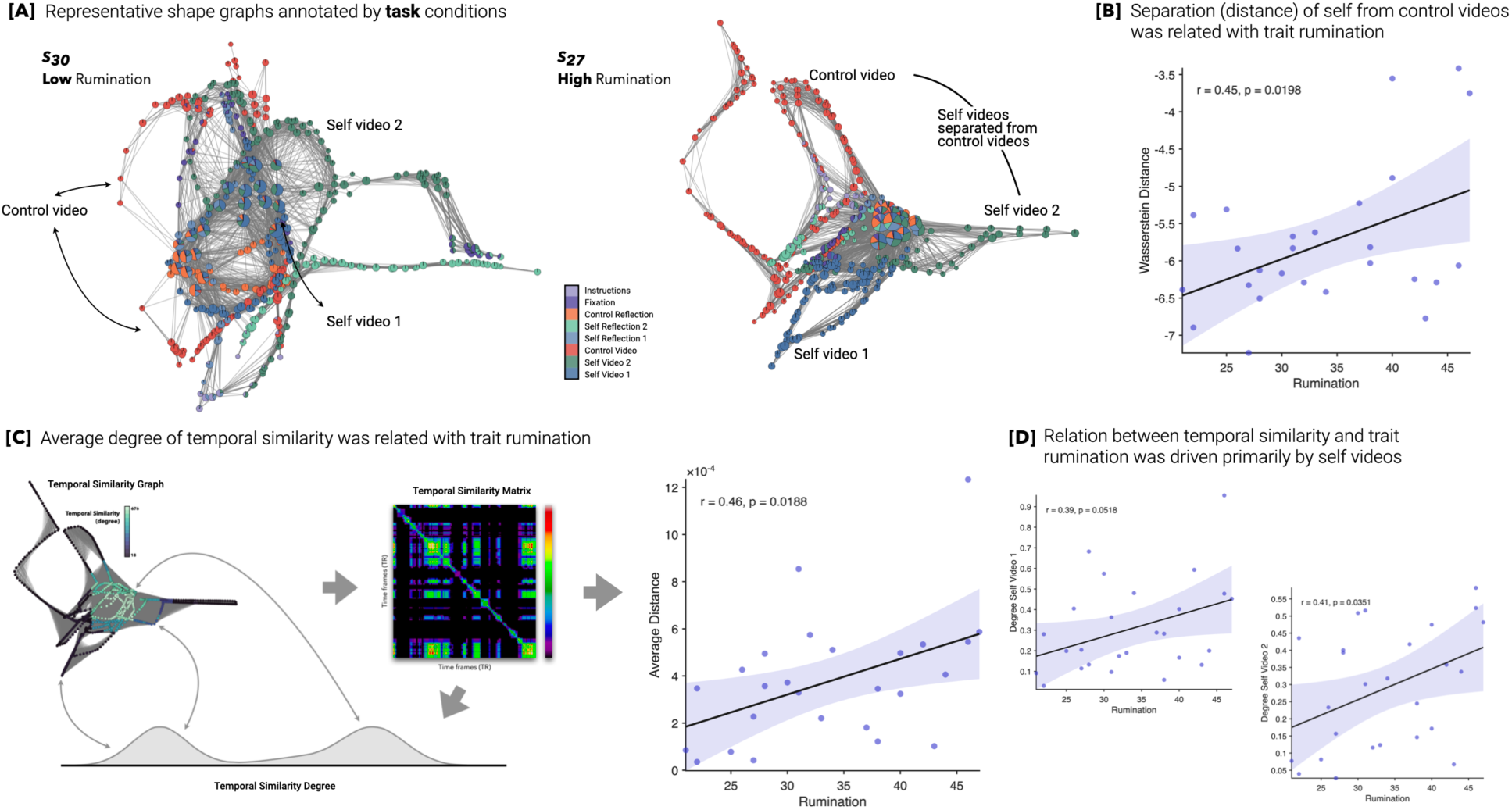
Mapper-derived brain-state transitions reveal altered dynamics during self-viewing in high trait ruminators. (**A**) Example Mapper graphs from two participants—one with low and one with high trait rumination—annotated by task conditions. In the high ruminator, brain states evoked by *self videos* are more topologically separated from those evoked by *control videos*, suggesting altered transitions between externally and internally focused cognitive states. (**B**) Across participants, the Wasserstein distance between self and control video-evoked state distributions increased with trait rumination (r = 0.45, *p* = 0.0198). (**C**) Higher trait rumination was also associated with a greater *average degree* in the temporal similarity graph, reflecting stronger local similarity between nearby timepoints (r = 0.46, *p* = 0.0188). (**D**) This effect was primarily driven by increased temporal similarity during *self-viewing video* conditions (Self-Viewing Video 1: r = 0.39, *p* = 0.0518; Self -Viewing Video 2: r = 0.41, *p* = 0.0351). Together, these findings suggest that individuals with high rumination exhibit more rigid and segregated brain dynamics during self-relevant reflection.

As a global measure of Mapper graph structure across conditions, we first computed the Wasserstein distance between nodes corresponding to self-viewing and control video segments, within each participant’s Mapper graph (**Figure 7B**). This analysis quantifies how topologically distinct the brain state trajectories are between the two task conditions. Surprisingly, we found a positive correlation between Wasserstein distance and trait rumination (Pearson’s r = 0.45, p = 0.0198), indicating that individuals higher in rumination show greater topological separation between self-relevant and control videos. This suggests that high ruminators differentiate more strongly between internally focused and externally focused states, possibly reflecting heightened salience or engagement with self-relevant stimuli.

Next, we focused on a complementary node-level measure, temporal similarity degree, defined as the number of temporal neighbors each node is connected to. This metric reflects how often a given brain state re-occurs or co-varies with other timepoints, i.e., a higher degree indicates lower variability in moment-to-moment brain dynamics. We hypothesized that individuals with higher trait rumination would exhibit more rigid brain state trajectories, resulting in a higher average degree.

To test this, we computed each participant’s average degree across all timepoints and correlated it with their trait rumination scores. As shown in **Figure 7C**, we observed a significant positive correlation (Pearson’s r = 0.46, p = 0.0188), supporting the hypothesis that trait rumination is associated with reduced variability or repetitiveness in brain dynamics during naturalistic reflection. This aligns with prior behavioral and clinical characterizations of rumination as a perseverative cognitive style.

To further localize this effect to specific task contexts, we repeated the degree analysis separately for each video condition, computing the average degree only over timepoints associated with each video condition. As shown in **Figure 7D**, we found that the temporal similarity degree during the self-viewing videos was strongly correlated with trait rumination (Self Video 1: r = 0.39, p = 0.0518; Self Video 2: r = 0.41, p = 0.0351). In contrast, the control video showed a nonsignificant association (r = 0.34, p = 0.09).

Together, these results highlight two complementary neural signatures of trait rumination: (1) increased topological differentiation of brain states across task contexts (Wasserstein analysis), and (2) decreased moment-to-moment variability in brain state transitions (degree analysis), especially during self-relevant conditions. The Mapper-based framework offers a powerful lens to uncover such dynamic, individualized neural patterns underlying complex cognitive traits like rumination.

### 3.4 Parameter Perturbation Analysis to Evaluate Stability of Mapper-derived Results

Topological data analysis methods such as Mapper rely on user-defined parameters (e.g., resolution, gain) that influence the granularity and connectivity of the resulting graph structures (Saggar et al., 2018). To assess the stability of our findings, we conducted a parameter perturbation analysis, systematically varying Mapper resolution and gain over 25 combinations (resolution = 10:30, steps of 5; and gain = 50:70, steps of 5).

#### Visual inspection of graph stability

Despite the expected variation in graph size and density across parameter settings, the overall topological properties of each participant’s Mapper graph remained visually stable. As shown in Supplementary **Figure S1**, participants with high trait rumination (e.g., Subj 27) consistently showed denser, more recurrent trajectories, particularly during self-relevant conditions, compared to low ruminators (e.g., Subj 30), whose graphs exhibited more dispersed and differentiated temporal patterns. This qualitative consistency across parameter space suggests that core features of Mapper graphs are robust to moderate perturbations in resolution and gain.

#### Stability of behavior-Mapping associations

To quantitatively assess robustness, we computed correlations between Mapper-derived metrics (e.g., mean temporal degree, Wasserstein distance) and trait rumination scores at each of the 25 parameter settings. For each behavioral association, we summarized the distribution of correlation coefficients by reporting the mean Pearson’s r and 95% confidence interval (CI) across the parameter grid.

For the mean temporal similarity degree, the average correlation across parameter combinations was r = 0.270, with a 95% CI = [0.075, 0.465], indicating a robust and statistically reliable positive association with trait rumination. This suggests that the observed effect of increased temporal rigidity in high ruminators is not a fragile artifact of specific Mapper parameter choices. In contrast, the correlation between Wasserstein distance, a measure of topological separation between self-related and control timepoints within the Mapper graph, and trait rumination was more variable across parameters. The average correlation was r = 0.060, with a 95% CI = [–0.129, 0.250], indicating non-significant and unreliable associations when parameters are perturbed.

Together, these results support the robustness of the Mapper-derived temporal similarity degree metric in capturing meaningful individual differences in rumination traits. In contrast, the correlation between Wasserstein distance and trait rumination did not survive parameter perturbation, suggesting this association is less reliable under varying topological constructions. However, it is worth noting that this may reflect limited statistical power or subtle effect sizes rather than the absence of a meaningful relationship. We include the Wasserstein result here as a potentially promising direction for future studies with larger sample sizes and improved sensitivity, particularly given its conceptual relevance in quantifying topological separation between cognitive states.

### 3.5 Anchoring Temporal Similarity to Brain Anatomy

To investigate the neurophysiological substrates underlying the topological signatures of brain dynamics observed in Mapper graphs, we performed a brain-wide analysis linking the degree of temporal similarity to regional brain activity.

We began by converting each participant’s Mapper-derived graph into a time-resolved format that preserved the degree of temporal co-occurrence for each fMRI timeframe (i.e., TR). This transformation allowed us to map the abstract topological property, temporal similarity degree, back onto brain anatomy. Specifically, for each task condition, we computed a weighted brain map in which each timepoint’s BOLD signal was modulated by its corresponding degree value (**Figure 8A**). These degree-weighted time series were then averaged within each task condition (e.g., self-viewing videos, control video) to create participant-specific explanatory variables for a within-subject GLM.

**Figure 8.**
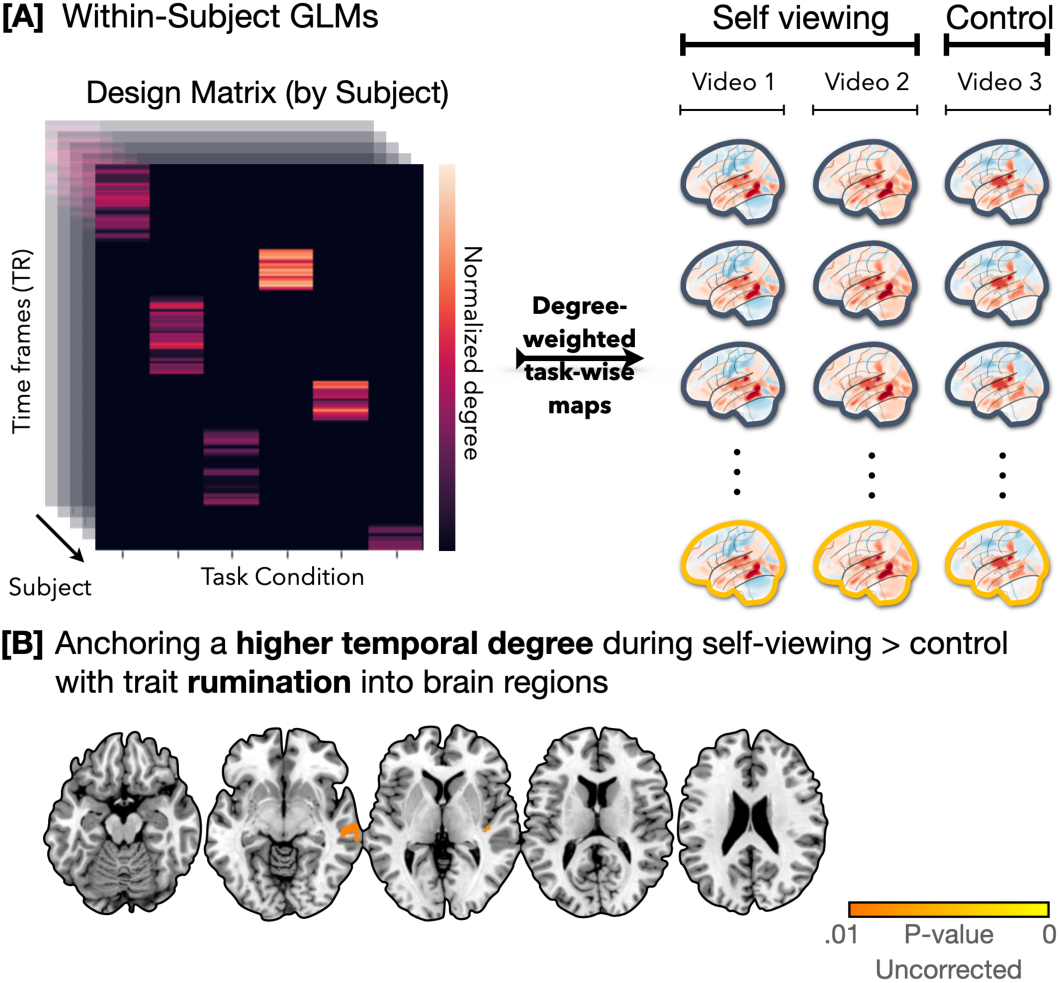
Anchoring Mapper-derived temporal similarity to brain anatomy using GLM analysis. (**A**) For each participant, Mapper graphs were decomposed into time-resolved vectors representing the degree of temporal similarity (i.e., the number of neighboring timepoints) for each fMRI volume. These vectors were normalized and aligned with task regressors to generate degree-weighted design matrices. These were then applied to the corresponding BOLD time series to compute task-wise brain maps (e.g., Self-viewing Videos 1 & 2, Control Video 3), yielding individualized representations of brain activity modulated by topological dynamics. (**B**) At the group level, we tested whether greater degree during self-viewing versus control was associated with higher trait rumination, using randomise with TFCE. While no clusters survived correction, uncorrected results at p < 0.01 revealed suggestive engagement in the right middle temporal gyrus and posterior insula, regions previously linked to interoception and social-emotional processing. These exploratory findings demonstrate how temporal dynamics derived from Mapper can be anchored to anatomical substrates of cognitive-affective traits.

At the group level, we then examined whether the relationship between degree-weighted BOLD activation and trait rumination varied across task conditions. We focused on the contrast:

(Self-viewing > Control video) × Trait Rumination and used FSL’s randomise with Threshold-Free Cluster Enhancement (TFCE) to identify clusters showing significant associations.

No clusters survived TFCE-corrected thresholds. However, when visualizing uncorrected results at a stricter threshold (p < 0.01, uncorrected), we observed suggestive activity in the right middle temporal gyrus and posterior insula, regions implicated in social cognition, interoception, and affective salience (**Figure 8B**). While exploratory, these findings provide a neuroanatomical anchor for the topological metric of temporal similarity degree and suggest that individuals with high trait rumination may exhibit distinctive activation patterns in specific cortical areas during naturalistic self-relevant processing.

## 4. Discussion

The tendency towards repetitive negative thought about one’s own affective state (rumination) has adverse effects on well-being and cognition. To study the neural bases associated with trait rumination, we characterized whole-brain fMRI dynamics while individuals watched and reflected on recordings of their own previous social interactions. We used the Rumination-Reflection Questionnaire to measure trait rumination. In a follow-up session, fMRI data were collected while individuals watched multiple 3-minute video recordings of their own team’s brainstorming session (as well as one showing a control team) and reflected on the team’s performance for 90 seconds following each video. We perform traditional general linear modeling (GLM) and inter-subject correlation (ISC) to further validate our novel naturalistic fMRI paradigm. We then extend the Mapper algorithm from TDA to create graphical representations of changes in whole-brain activity across the video watching and reflection tasks, and we introduce new ways to quantify these representations in a behaviorally relevant manner. Lastly, we anchor Mapper findings to the brain’s anatomy.

### Task-based GLM analysis

revealed that viewing self-related videos, compared to viewing others, engaged a distributed set of regions implicated in self-referential and social evaluative processing. Specifically, we observed significantly increased activity in the anterior cingulate cortex (ACC), supplementary motor area (SMA), bilateral middle and inferior temporal gyri, and fusiform cortex. These results align with prior findings showing that the ACC plays a key role in performance monitoring and self-evaluation, particularly in emotionally salient contexts (Denny et al., 2012; Etkin et al., 2011). The fusiform and inferior temporal regions are traditionally associated with processing socially relevant visual features, including faces and identity cues (Hoffman & Haxby, 2000; Kanwisher & Yovel, 2006), which is consistent with participants viewing videos of themselves. The involvement of the middle temporal gyrus may reflect semantic and autobiographical processing, which has also been implicated in narrative-based self-reflection (Spreng & Grady, 2010)

These findings support the construct validity of our naturalistic task design. Despite using unscripted, real-world stimuli, our paradigm elicited robust activation in regions commonly associated with self-related cognition and social context appraisal. Importantly, the anterior midline structures identified, such as the dACC, overlap with the default mode network (DMN), a system strongly linked to internal mentation, including self-focused thought and rumination (Andrews-Hanna et al., 2014). Thus, the increased recruitment of these areas during self-viewing suggests that our paradigm successfully taps into the neurocognitive mechanisms of self-referential processing under ecologically valid conditions. Notably, the engagement of both perceptual and introspective systems mirrors the dual demands of the task, i.e., watching oneself interact while simultaneously evaluating one’s social performance, which may be particularly relevant for individuals prone to rumination.

### ISC analysis

revealed increased neural synchrony across individuals during self-viewing compared to control viewing, specifically localized to the right cerebellar lobules I–IV. Although the cerebellum is traditionally associated with motor control, an expanding body of research underscores its involvement in higher-order cognitive and affective processes, including social cognition and self-referential processing (Buckner et al., 2011; Van Overwalle et al., 2020). The cerebellum’s role in timing and predictive modeling may be especially important in dynamic, naturalistic contexts such as watching oneself in a prior social interaction, where it may help simulate and evaluate interpersonal behavior.

Moreover, the right anterior cerebellum, particularly lobules I–IV, has been implicated in sensorimotor integration and embodied self-awareness functions that may be recruited when individuals evaluate their own bodily expressions, tone, and gestures in a video of themselves. The observed synchrony suggests that participants engaged similar cerebellar computations while processing self-relevant social stimuli, possibly reflecting a shared sensorimotor or affective evaluation schema. Importantly, this pattern emerged despite self-videos varying across teams while the control video remained identical, a design that would typically favor greater stimulus-driven synchrony during control viewing. The reversed pattern observed here suggests that self-relevance elicited common cerebellar processing that was not simply driven by low-level perceptual features.

That this cluster was the only significant ISC finding also underscores the specificity of cerebellar engagement with subtle, self-relevant cues, in contrast to cortical synchrony, which may be more variable or diluted across individuals in complex, naturalistic tasks. The use of participant-specific ISC scores allowed us to preserve this variability and capture heterogeneity in neural alignment, supporting recent calls to move beyond traditional group-averaged ISC methods when studying individual differences (Finn et al., 2018, 2020).

### Mapper-based analysis

revealed a novel link between trait rumination and reduced variability in brain dynamics during a naturalistic self-reflection task. This topological characterization aligns with and extends prior evidence suggesting that *mental inflexibility* is a hallmark of rumination and related mood disorders. Rumination is often conceptualized as a maladaptive form of cognitive perseveration or mental entrapment, characterized by repetitive, internally focused thought loops (Nolen-Hoeksema et al., 2008; Watkins, 2008). Several studies using traditional functional connectivity approaches have reported decreased variability or flexibility of brain networks in high ruminators (Demirtas et al., 2016; Gao et al., 2023; Hamilton et al., 2011; Li et al., 2022; Peterson et al., 2025).

In particular, prior work has shown that state rigidity, or reduced switching between distinct brain network configurations over time, is associated with negative affect and poor mental health outcomes (Damaraju et al., 2014; Vidaurre et al., 2017). Our findings, using Mapper to capture these dynamics at the individual level, provide a geometry-preserving, model-free framework to visualize and quantify such rigidity. The positive association between trait rumination and temporal similarity (degree) suggests that high ruminators exhibit less differentiated brain states over time, which is consistent with a “sticky” or inflexible brain that repeatedly samples similar configurations (Demirtas et al., 2016; Gao et al., 2023; Hamilton et al., 2011; Li et al., 2022; Peterson et al., 2025).

Moreover, the task-specific nature of the Mapper findings, i.e., significant relationships between rumination and temporal similarity during self-viewing but not during control viewing, highlights the importance of self-relevant contexts in eliciting rigid brain dynamics. This adds a novel dimension to prior work, which has primarily examined rumination-related neural rigidity during resting-state conditions (e.g., (Andrews-Hanna et al., 2014; H.-X. Zhou et al., 2020)). By contrast, our study leverages an ecologically valid, naturalistic paradigm that explicitly engages self-evaluative processes through first-person video stimuli. This task-evoked context may more faithfully approximate the real-world scenarios in which rumination typically unfolds, and demonstrates for the first time that temporal inflexibility in large-scale brain activity extends beyond rest and manifests during active self-appraisal.

Importantly, the Mapper framework enables us to quantify this inflexibility at the level of brain-state transitions, independent of stimulus timing, a crucial feature for studying individual responses to naturalistic, self-relevant stimuli. Prior studies examining rumination and attractor-like dynamics have largely relied on resting-state fMRI, demonstrating reduced metastability and lingering within narrow brain activity patterns in high ruminators (Rolls, 2021). However, our study is among the first to elicit ruminative tendencies through a naturalistic task paradigm and show that reduced brain dynamics (i.e., lower transition diversity) are significantly associated with trait rumination during this self-referential viewing. Notably, conventional analyses, including task-based activation (GLM) and inter-subject synchrony (ISC), identified condition-sensitive clusters, but these did not correlate with individual rumination scores. This underscores the advantage of Mapper-based representations in capturing fine-grained, individualized neural dynamics that may be obscured in group-level, spatially constrained analyses.

These findings align with emerging theoretical models, such as those proposed by Zhou et al. (2025), who used network control theory and task-based fMRI to show that individuals with depression or anxiety exhibit lower control energy in key nodes (e.g., ACC, lateral PFC) when attempting to switch away from perseverative thought (D. Zhou et al., 2025). They interpret this as entrenchment in low-energy attractor states, akin to how a ball settles in the bottom of a deep well. Our Mapper-based topological characterization complements this model by offering an unsupervised, data-driven quantification of such “stickiness” in brain activity trajectories, with lower transition diversity reflecting a similar tendency to remain trapped within a small number of attractor basins.

By further mapping temporal similarity onto anatomy, we observed convergence in the right middle temporal gyrus and posterior insula (uncorrected). These regions are implicated in autobiographical memory (Spreng et al., 2009) and interoceptive awareness (Craig, 2009), respectively. This suggests that high-ruminating individuals may be more prone to engaging in viscerally grounded, self-referential simulations when viewing themselves, and that these simulations exhibit reduced neural flexibility compared to those with lower trait rumination.

### Robustness and Limitations of Topological Findings

While our primary Mapper-derived measure, i.e., temporal similarity degree, demonstrated a robust association with trait rumination across multiple parameter settings, not all TDA-derived metrics exhibited this level of consistency. Specifically, we observed a positive correlation between Wasserstein distance (quantifying topological separation between self-viewing and control brain states) and trait rumination. This suggested a potential signature of heightened self-related segregation in high ruminators. However, this effect did not survive parameter perturbation analysis, indicating reduced reliability. Given the conceptual relevance of this measure and the limited sample size of the current study, we interpret this null result cautiously and include it as a promising exploratory direction for future work. Larger and statistically powered samples may be better positioned to detect subtle yet meaningful topological distinctions in how self-referential and neutral stimuli are processed.

Altogether, by implementing a comprehensive approach that combined traditional activation (GLM), synchrony (ISC), and dynamic TDA (Mapper) analyses, our study offers a convergent view of how trait rumination manifests in brain function during self-relevant naturalistic tasks. The GLM results showing elevated engagement of the anterior cingulate, supplementary motor area, bilateral temporal gyri, and fusiform cortex during self-viewing align with long-standing findings that self-referential cognition recruits midline and lateral brain systems (Denny et al., 2012; Etkin et al., 2011; Kanwisher & Yovel, 2006). The ISC findings then add a layer of interpersonal synchrony: during self-viewing, participants showed increased shared activity in the right cerebellar lobules I–IV, suggesting common embodied or sensorimotor simulation processes across subjects (Buckner et al., 2011; Van Overwalle et al., 2020). Finally, the Mapper results extend these results into the temporal-dynamic domain: individuals with higher trait rumination exhibited brain state trajectories with higher average temporal similarity degree, that is, reduced variability and more recurrent brain states over time. This dynamic signature is consistent with literature linking rumination to cognitive rigidity, reduced network flexibility, and altered dynamic functional connectivity (Zhang et al., 2022; Kim et al., 2023).

Importantly, the convergence of findings across methods suggests that rumination may not only be about which brain regions are active (GLM) or how similar participants are to one another (ISC), but also how the brain moves from state to state over time (Mapper). In particular, while GLM and ISC focus on averaged or group-level features of brain response, Mapper preserved individual variability in dynamic transitions, thus making it highly suited for detecting trait-relevant differences in brain dynamics. This is in line with recent calls to examine brain dynamics (rather than static connectivity or activation) as markers of rumination and psychopathology risk (Vidaurre et al., 2017; Damaraju et al., 2014). Furthermore, our finding that the trait-rumination correlation with degree was strongest during self-viewing (vs. control) reinforces perspectives that self-relevant or autobiographical contexts amplify individual differences in brain function (Andrews-Hanna et al., 2014; Zhou et al., 2020).

### Limitations and future work

Several limitations of the current study should be acknowledged. First, the sample size was modest, which may limit statistical power, particularly for higher-order interactions or for detecting small effect sizes in ISC or Mapper-based metrics. While we observed robust associations between Mapper-derived brain dynamics and trait rumination, these findings should be replicated in larger, more diverse samples to confirm their generalizability.

Second, although the naturalistic self-reflection paradigm provides high ecological validity, it introduces challenges in experimental control. Variability in participants’ engagement, memory recall, and interpretation of their own behavior may have contributed to unmeasured heterogeneity. While this variability is arguably a feature rather than a flaw, given the goal of capturing real-world self-evaluative processes, future studies might incorporate online behavioral or physiological measures (e.g., eye tracking, skin conductance) to index engagement and help interpret brain responses.

Third, while our Mapper analysis offers a powerful method for quantifying individual brain dynamics, its interpretability depends on several parameter choices (e.g., filter functions, cover size, clustering method). We partially mitigated this concern by choosing parameters based on heuristics informed by prior studies and running parameter perturbation analysis for robustness.

Fourth, our analysis of anatomical correlates of Mapper-derived metrics revealed uncorrected effects in regions such as the middle temporal gyrus and posterior insula. While suggestive, these effects did not survive correction for multiple comparisons, likely due to sample size. Replication in larger datasets is needed to clarify whether these regions consistently anchor the temporal rigidity observed in high ruminators.

Lastly, the cross-sectional design of the study limits inferences about causality. It remains unclear whether reduced dynamic variability is a cause or consequence of ruminative tendencies, or whether both reflect underlying trait-level differences in cognitive control, self-focus, or emotional regulation. Longitudinal designs, or studies that combine neurofeedback or intervention approaches (e.g., mindfulness, cognitive reappraisal training), may help clarify the directionality of these effects and their malleability.

### Conclusions

Our study demonstrates that naturalistic, self-relevant paradigms, when combined with modern analytical tools such as Mapper and ISC, can reveal novel individual-level insights into the neurocognitive underpinnings of rumination. We show that trait rumination is associated with reduced temporal variability in brain dynamics, as captured by Mapper-derived graphs, particularly during self-relevant conditions. These dynamic rigidity patterns align with prior theories positing rumination as a repetitive and inflexible mode of internal mentation. Notably, the integration of traditional GLM and ISC findings with topological dynamics strengthens the robustness of these interpretations.

Together, our results provide a multidimensional portrait of how the ruminative mind manifests in both static and dynamic neural processes during ecologically valid self-reflection. By grounding temporal graph features in established anatomical networks and capturing behaviorally relevant variability across individuals, this work contributes toward a mechanistic understanding of maladaptive self-focus. More broadly, these findings suggest that dynamic neural rigidity may serve as a transdiagnostic marker of maladaptive self-focus, opening avenues for precision interventions targeting cognitive flexibility.

## Acknowledgements

This work was supported by an HPI-Stanford Hasso Plattner Design Thinking Research Program (HPDTRP) award and an NIH Director’s New Innovator award (MH-119735) to M.S.

## Data and Code availability

Mapper is available here https://github.com/braindynamicslab/dyneusr. Data is available upon request.

## Supplementary Information

**Figure S1:**
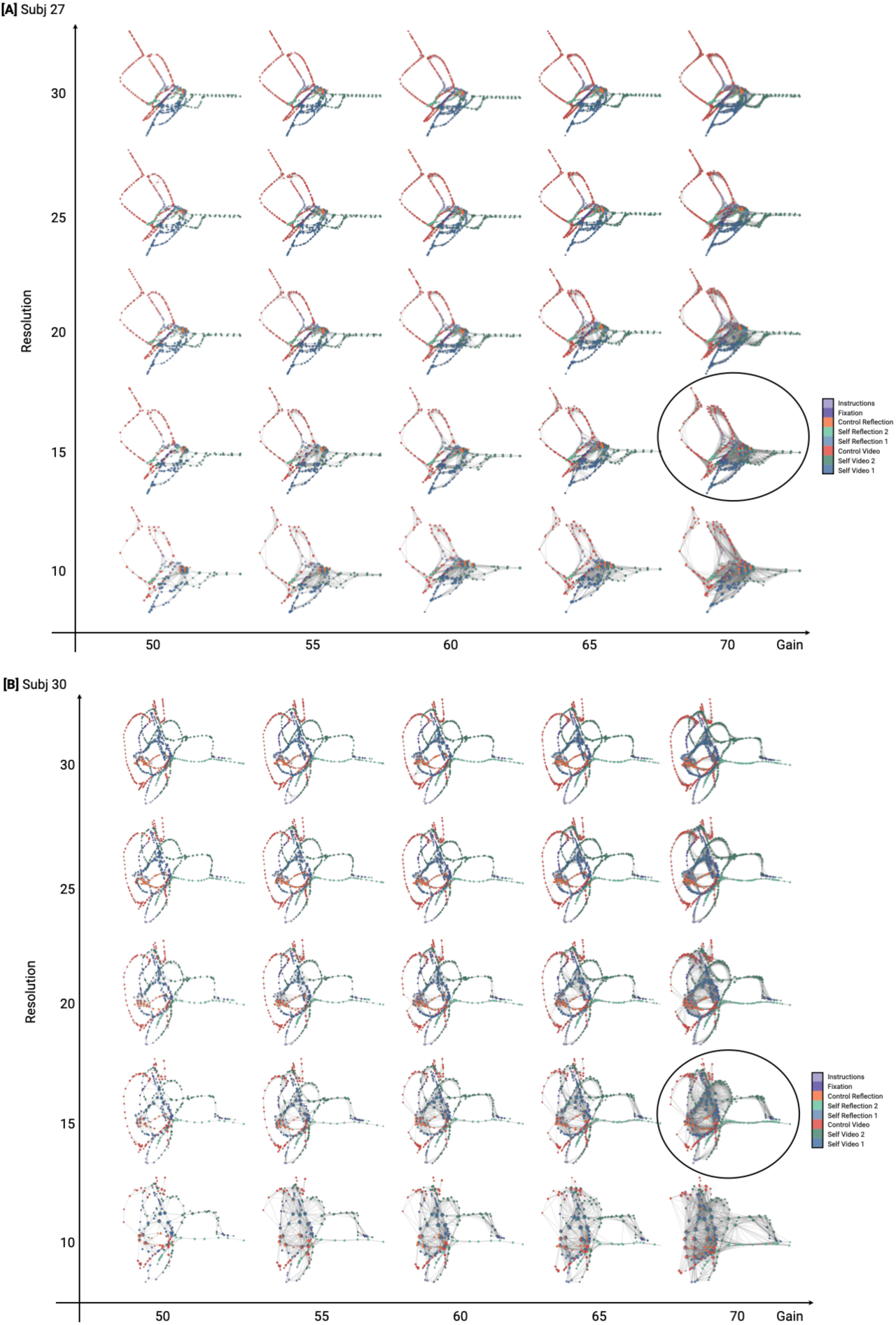
Parameter perturbation for two representative participants. A wide variety of Mapper parameters did not yield qualitative differences in the overall properties of Mapper graphs. Circled parameters were used in the current work (resolution =15, gain = 70%).

